# Genomic Surveillance Reveals Global Spread of Macrolide-Resistant *Bordetella pertussis* Linked to Vaccine Changes

**DOI:** 10.1101/2025.07.16.665123

**Authors:** Zhen Xu, Zhuoying Huang, Lingyue Yuan, Huanyu Wu, Xin Chen, Min Chen, Yuan Zhuang, Jun Feng

## Abstract

The resurgence of whooping cough in regions utilizing acellular pertussis vaccines underscopes emerging public health challenges. Here, we characterized 178 *Bordetella pertussis* isolates collected from patients across all age groups in Shanghai (2018-2024) to assess genomic evolution and antibiotic susceptibility. Macrolide resistance to erythromycin, azithromycin, clarithromycin and clindamycin escalated from ≤50% (pre-2020) to nearly 100% (post-2020), mechanistically linked to the 23S rRNA A2047G mutation. Genome-based analysis identified a genotype MT28-*ptxP3*-MRBP rapidlly dominated post-2020, exhibiting significantly higher prevalence in adults versus than age groups. Phylogenetic analysis of 178 Shanghai and 1596 global genomes revealed two major lineages corresponding to *ptxP1* and *ptxP3* alleles. MT28-*ptxP3*-MRBP cluster was identified in France, Japan and the United States in 2024, indicating potential cross-border dissemination. These findings advocate for intergrated surveillance spanning all ages and international borders to contain the global spread of macrolide-resistant *Bordetella pertussis*.

**Highlights:**

1. After 2020, MT28-*ptxP3*-MRBP lineage rapidly dominated, comprising 61.7% of isolates.
2. MT28-*ptxP3*-MRBP exhibits a significant transmission advantage among older individuals.
3. The primary affected group shifted from ≤36 months (pre-2020) to 37 months–18 years (post-2020).
4. Macrolide resistance rose from ≤50% pre-2020 to nearly 100% post-2020, with all resistant isolates carrying the A2047G mutation.

## Introduction

Whooping cough, an acute respiratory infectious disease caused primarily by *Bordetella pertussis* (*B. pertussis*), is characterized by high transmissibility and poses a severe threat to young infants ^[1]^. Over the past two decades, countries with high coverage of acellular pertussis (aP) vaccination—including the United States, France, and New Zealand—have witnessed a resurgence of whooping cough from historically low incidence in the early 21st century, with regional outbreaks defining the phenomenon of “pertussis resurgence”^[2-4]^. Since 2012, China has exclusively used aP vaccines ^[5]^, maintaining a coverage rate exceeding 97% ^[6]^. Paradoxically, reported cases surged from 2016, reaching over 30 000 in 2019—the highest levels since the late 1980s ^[6, 7]^.

Notebly, *B. pertussis* can infect or reinfect individuals across all age groups ^[8]^, yet the true disease burden in adults remain significantly underestimated ^[9, 10]^. Current studies on antibiotic susceptibility and genomic evolution of Chinese *B. pertussis* isolates predominantly focus on pediatric populations^[11-13]^,, with limited systematic investigations across other age demographics. This gap hinders comprehensive assessment of resistance dynamics and transmission risk.

Here, we leveraged Shanghai’s active surveillance system to characterize 178 *B. pertussis* isolates from patients of diverse age groups (2018–2024). Through antibiotic susceptibility testing and whole-genome analyses, we evaluated trends in macrolide resistance, and characterize molecular evolutionary features, and, integrate epidemiological data to explore age-related distribution patterns and potential risks of international dissemination.

We observe a dramatic surge in macrolide resistance, with rates escalating from ≤ 50% (pre-2020) to nearly 100% (post-2020). All resistant isolates harbored the 23S rRNA A2047G mutation, a hallmark of macrolide resistance in *B. pertussis*. Multilocus variable-number tandem repeat analysis (MLVA) and vaccine antigen genotyping revealed rapid expansion of the MT28-*ptxP3* lineage of macrolide-resistant *B. pertussis* (MRBP) after 2020, with a significantly higher prevalence in adults compared to other age groups; Global phylogenetic analysis further demonstrated the detection of this lineage in France, Japan, and the United States in 2024, indicating a potential risk of cross-border transmission. These findings underscore the critical need for continuous, age-stratified surveillance of *B. pertussis* infections. The rapid emergence and international dissemination of MRBP highlight the urgency of enhancing global collaborative efforts to address this evolving public health challenge.

## Materials and methods

### Bacterial isolates

According to the Shanghai pertussis active‐surveillance protocol, from 2018 through 2023 six sentinel hospitals were included: Hospital A in Minhang District, Community Health Service Center A in Putuo District, Medical Center A and Hospital B in Pudong New Area, and Hospital C and Community Health Service Center B in Songjiang District. In 2024, surveillance was expanded to ten sites with the addition of Hospital D in Xuhui District, Hospital E in Minhang District, and Hospital F plus Community Health Service Center C in Qingpu District. For all patients meeting the case definition, nasopharyngeal swabs were collected and immediately transported to the Bacterial Testing Laboratory at Shanghai CDC. Upon receipt, specimens were plated onto charcoal‐selective agar (Qingdao Zhongchuang Huike Biotechnology Co., Ltd., China) and incubated at 36 °C in 5% CO_2_ for 3-7 days. After *B. pertussis*– specific nucleic acid fragments were confirmed using a commercial PCR kit (Jiangsu Bioperfectus Technologies Co., Ltd., China), single colonies were subcultured onto charcoal agar and stored in milk‐based preservation tubes at –80 °C for future analyses. This study was approved by the Ethics Committee of Shanghai CDC (Approval No. KY-2025-15).

### Antimicrobial sensitivity tests

Bacterial suspensions from milk‐based preservation tubes were inoculated onto charcoal agar plates and incubated at 36 °C in 5% CO_2_ for 72 h to revive *B. pertussis*. A 0.5 McFarland standardized suspension was prepared in API 0.85% NaCl solution (bioMérieux, France) and uniformly plated onto charcoal agar. E-test strips (Liofilchem, Italy) for erythromycin (ERY), azithromycin (AZM), clarithromycin (CLR), and clindamycin (CLI) were applied, and plates were incubated at 36 °C in 5% CO_2_ for 72 h to determine minimum inhibitory concentrations (MICs). As neither the Clinical and Laboratory Standards Institute (CLSI) nor the European Committee on Antimicrobial Susceptibility Testing (EUCAST) has defined specific breakpoints for *B. pertussis*, an MIC ≥ 32 mg/L was considered resistant to all four antibiotics^[14]^.*B. pertussis* ATCC 9797 was used as the quality control strain.

### Whole-genome data sources, assembly and screening

Genomic DNA from revived *B. pertussis* isolates was extracted using the QIAamp PowerFecal Pro DNA Kit (QIAGEN, Germany) according to the manufacturer’s instructions. DNA purity and concentration were evaluated using a NanoDrop spectrophotometer (Thermo Fisher Scientific, USA) and a Qubit 4.0 Fluorometer (Thermo Fisher Scientific, USA), respectively. Samples meeting quality criteria were subjected to paired-end 150 bp (PE150) sequencing on both the DNBSEQ-T7 platform (MGI, China) and the NextSeq 2000 platform (Illumina, USA). In addition, publicly available *B. pertussis* genomes deposited in NCBI from January 2016 through May 2025 were retrieved for comparative analysis.

Raw FASTQ reads were quality-controlled with fastp v0.23.4 and de novo assembled using SPAdes v3.15.5, retaining contigs > 1000 bp. Both newly assembled and downloaded FASTA genome sequences were taxonomically classified with Kraken2 v2.1.3 and assessed for completeness and contamination using CheckM2 v1.0.2. Only assemblies identified as *B. pertussis* with ≥ 99.9% completeness and ≤ 1.5% contamination were included in downstream analyses. In total, 1774 *B. pertussis* genomes were analyzed, including 178 isolates sequenced in this study and 1596 isolates retrieved from NCBI databases (see Supplementary Table 1 for epidemiological information on the downloaded isolates).

**Table 1.** Epidemiological characteristics of *B. pertussis* isolates, Shanghai, 2018-2024

| Year | Total | 2018 | 2019 | 2021 | 2022 | 2023 | 2024 |
| --- | --- | --- | --- | --- | --- | --- | --- |
| Culture-Confirmed | 178 | 24 | 26 | 35 | 27 | 11 | 55 |
| Gender |  |  |  |  |  |  |  |
| Male | 93 | 14 | 11 | 19 | 17 | 7 | 25 |
| Female | 85 | 10 | 15 | 16 | 10 | 4 | 30 |
| Age |  |  |  |  |  |  |  |
| < 3 months | 26 | 9 (37.50%) | 7 (26.92%) | 2 (5.71%) | 3 (11.11%) | 2 (18.18%) | 3 (5.45%) |
| 3-6 months | 22 | 5 (20.83%) | 7 (26.92%) | 7 (20.00%) | 2 (7.41%) | 0 | 1 (1.82%) |
| 7-36 months | 10 | 5 (20.83%) | 2 (7.69%) | 2 (5.71%) | 0 | 0 | 1 (1.82%) |
| 37 months-6 years | 32 | 0 | 1 (3.85%) | 11 (31.43%) | 7 (25.93%) | 2 (18.18%) | 11 (20.00%) |
| 7-18 years | 41 | 0 | 0 | 1 (2.86%) | 10 (37.04%) | 3 (27.27%) | 27 (49.09%) |
| 19-40 years | 19 | 4 (16.67%) | 4 (15.38%) | 10 (28.57%) | 0 | 1 (9.09%) | 0 |
| >40 years | 28 | 1 (4.17%) | 5 (19.23%) | 2 (5.71%) | 5 (18.52%) | 3 (27.27%) | 12 (21.82%) |

### 23S rRNA A2047G mutation detection

The A2047G mutation in the *B. pertussis* 23S rRNA gene was detected by two complementary approaches. First, assembled genomes were aligned to the Tohama I reference (GenBank accession GCA_000195715.1) using nucmer v3.1 to call nucleotide variants. Second, 23S rRNA loci were typed by querying the predefined alleles in the BIGSdb-Pasteur database; isolates classified as allele “13” were designated as harboring the resistance mutation^[15]^.

### Multiple locus variable-number tandem repeat (VNTR) analysis (MLVA)

MLVA was performed using the wgsMLVA pipeline as previously described by Weigand et al.^[16]^, and *B. pertussis* isolates were typed according to the five‐locus VNTR scheme (VNTR1, VNTR3a/VNTR3b, VNTR4, VNTR5, and VNTR6) proposed by Schouls et al ^[17]^.

### MLST and vaccine antigen genotyping

Isolates were typed by MLST using the scheme established in the BIGSdb-Pasteur database. Key vaccine antigen loci—including *ptxP, ptxA, ptxC, fhaB2400_5550, prn, fim2, fim3, and tcfA*—were then extracted from the database definitions to characterize the vaccine antigen genotype of each isolate. Because the full-length *fhaB* gene (∼10,773 bp) is often fragmented during genome assembly, the analysis was restricted to the *fhaB2400_5550* fragment as defined in the BIGSdb-Pasteur database^[15]^. Phylogenetic Analysis

Using the Tohama I reference genome (GenBank accession GCA_000195715.1), raw reads were aligned with Snippy v4.6.0 using default parameters, and recombinant regions were filtered out with Gubbins v2.4.1. The resulting core SNP alignment was used to infer a maximum‐likelihood phylogeny in IQ‐TREE v2.3.6 with automated model selection (−m MFP). Branch support was assessed by 1,000 ultrafast bootstrap replicates (−B 1000) and 1,000 SH-aLRT tests (−alrt 1000). All trees were visualized and edited on the Interactive Tree of Life (iTOL) web server (https://itol.embl.de/, accessed 15 June 2025). Genotypes not listed in the figure legends or not identified by these bioinformatic analyses were collectively designated as “Others.”

### Statistical analysis

To ensure consistency and accuracy amid incomplete age data for some pediatric cases, we defined the “parental” age group as 19–40 years and the “grandparental” age group as > 40 years. Strains were categorized into three temporal groups—pre-2020, 2020, and post-2020—based on prior studies^[11]^. All statistical analyses were performed using SPSS v25.0. Categorical variables were compared by X^2^ test or Fisher’s exact test, and a two-sided P-value < 0.01 was considered statistically significant.

## Results

### Epidemiological characteristics of the 178 culture-confirmed patients

A total of 2301 nasopharyngeal swabs yielded 178 *B. pertussis* isolates, with epidemiological characteristics detailed in Table 1. No isolates were recovered in 2020 due to low sampling, whereas the highest number of isolates occurred in 2024 (55/178, 30.90%). In the pre-2020 cohort, 70% of cases occurred in infants ≤ 36 months of age. In contrast, post-2020 cases predominantly occurred in school‐age children and adolescents (37 months-18 years, 52.17%). Two age-specific proportions differed significantly between periods (infants: χ^2^ = 44.31, p < 0.01; children/adolescents: χ^2^ = 43.74, P < 0.01), whereas cases in adults (≥ 19 years) showed no significantly change (χ^2^ = 0.091, p > 0.05).

### Antimicrobial susceptibility of *B. pertussis* and analysis of the A2047G resistance mutation

Of the 178 *B. pertussis* isolates, 152 (85.39%) exhibited MICs > 256 mg/L for all four tested antibiotics, while the remaining isolates showed MICs ≤ 1 mg/L (Fig. 1). After 2020, macrolide resistance rates surged from ≤ 50% to nearly 100%. Molecular assays confirmed that the 23S rRNA A2047G mutation exclusively in all resistant isolates, with no detection in susceptible isolates.

**Fig. 1.**
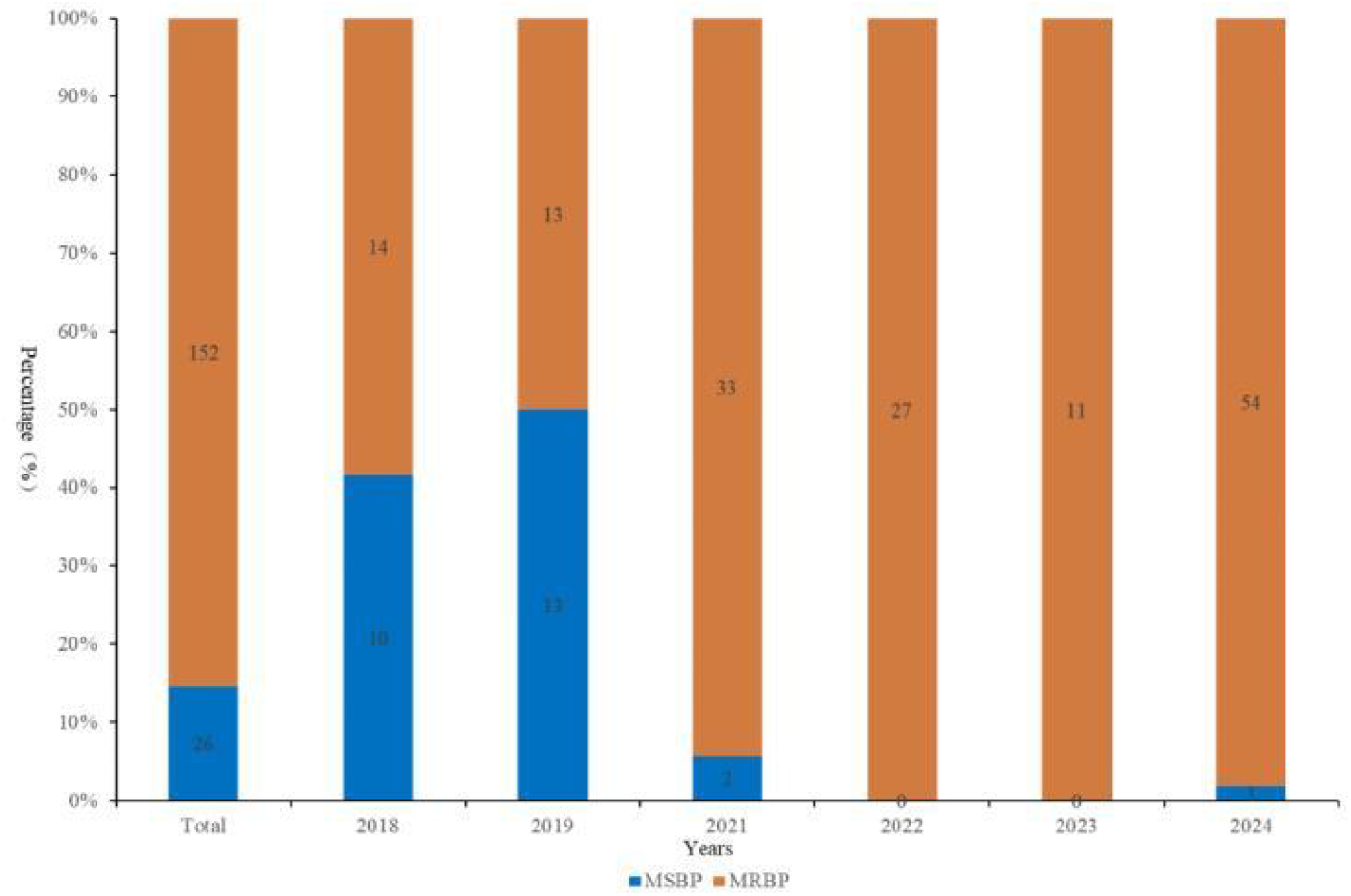
Macrolide susceptibility of *B. pertussis* isolates in Shanghai, 2018-2024. Stacked bars indicate the annual percentage of macrolide-sensitive *B. pertussis* (MSBP) and macrolide-resistant *B. pertussis* (MRBP) isolates, with the leftmost bar showing the overall (“Total”) distribution. Blue segments denote MSBP (MIC ≤ 1 mg/L) and orange segments denote MRBP (MIC ≥ 32 mg/L); the number within each segment indicates the count of isolates. The y-axis shows percentage of isolates, and the x-axis shows year of collection.

### Analysis of MLVA and multilocus sequence typing (MLST)

All isolates were assigned to ST2 by MLST. In addition, 17 distinct MLVA types (MTs) were identified, excluding nine isolates with unassignable MTs. The predominant types were MT28 (n = 87), MT195 (n = 26), MT60 (n = 22), and MT27 (n = 15) (Table 2). Notably, the prevalence of MT28 increased dramatically from 16% (8/50) pre-2020 to 61.17% (79/128) post-2020, paralleling a rise in macrolide– resistance from 0% (0/8) to 100% (79/79) with this lineage. In contrast, MT195 prevalence declined sharply from 48% (24/50) pre-2020 to 1.56% (2/128) post-2020. MT60, a post 2020 emergent type, accounted for 17.19% (22/128) of isolates, all of wihch were macrolide–resistant.

**Table 2.** Macrolide resistance of *B. pertussis* by MLVA type before and after 2020. * S: susceptible,R:resistance

|  | MT27 |  | MT28 |  | MT195 |  | MT60 |  | Others |  |
| --- | --- | --- | --- | --- | --- | --- | --- | --- | --- | --- |
|  | S | R | S | R | S | R | S | R | S | R |
| Pre-2020 | 12 (24%) | 0 | 8 (16%) | 0 | 0 | 24 (48%) | 0 | 0 | 3 (6%) | 3 (6%) |
| Post-2020 | 1 (0.78%) | 2 (1.56%) | 0 | 79 (61.72%) | 0 | 2 (1.56%) | 0 | 22 (17.19%) | 2 (1.56%) | 20 (15.63%) |

\* S: susceptible, R:resistance

### Vaccine antigen genotype analysis

Among the eight key vaccine antigen genes analyzed, only allele type 1 was detected for *ptxA, fim2*, and *fim3*, leading to their exclusion from further analysis. Six antigen genotype combinations were identified (Table 3). Pre-2020, the most prevalent genotypes were *ptxC2/prn2/ptxP3/fhaB2400_5550-1/tcfA2* (36%, 18/50) and *ptxC1/prn1/ptxP1/fhaB2400_5550-3/tcfA2* (48%, 24/50). Post-2020, *ptxC2/prn150/ptxP3/fhaB2400_5550-1/tcfA2* emerged as the predominant genotype (93.75%, 120/128). The temporal shift in antigen genotype composition from pre-to post-2020 can be divided into two main transitions: A change from a roughly equal distribution between *ptxC1/ptxP1/fhaB2400_5550-3* (46%, 23/50) and *ptxC2/ptxP3/fhaB2400_5550-1* (54%, 27/50) to near-exclusive predominance of *ptxC2/ptxP3/fhaB2400_5550-1* (96.88%, 124/128); A switch from predominantly *prn1* (48%, 24/50) and *prn2* (36%, 18/50) to overwhelmingly prn150 (94.53%, 121/128).

**Table 3.** Distribution of vaccine‐antigen genotype combinations, 2018-2024

|  | 2018 | 2019 | 2021 | 2022 | 2023 | 2024 |
| --- | --- | --- | --- | --- | --- | --- |
| *ptxC2/prn150/ptxP3/fhaB2400_5550-1/tcfA2* | 0 | 3 (11.54%) | 29 (82.86%) | 26 (96.30%) | 11 (100%) | 54 (98.18%) |
| *ptxC2/prn2/ptxP3/fhaB2400_5550-1/tcfA2* | 8 (33.33%) | 10 (38.46%) | 2 (5.71%) | 0 | 0 | 1 (1.82%) |
| *ptxC1/prn1/ptxP1/fhaB2400_5550-3/tcfA2* | 13 (54.17%) | 11 (42.31%) | 3 (8.57%) | 1 (3.7%) | 0 | 0 |
| *ptxC1/prn166/ptxP1/fhaB2400_5550-3/tcfA2* | 1 (4.17%) | 2 (7.69%) | 0 | 0 | 0 | 0 |
| *ptxC2/prn149/ptxP3/fhaB2400_5550-1/tcfA2* | 2 (8.33%) | 0 | 0 | 0 | 0 | 0 |
| *ptxC2/prn150/ptxP3/fhaB2400_5550-1/tcfA9* | 0 | 0 | 1 (2.86%) | 0 | 0 | 0 |

### Age‐specific differences in *B. pertussis* after 2020

Post-2020, the *ptxC2/prn150/ptxP3/fhaB2400_5550-1/tcfA2* vaccine antigen genotype dominated across all age groups (93.75%, 120/128), with no significant age-realted differences (χ^2^ = 10.48, p > 0.05) (Table 4). Concomitantly, macrolide resistance showed a similar age-indepentent pattern, with only three isolates retaining susceptiblity and no significantly variation in resistance rates (χ^2^ = 3.68, p > 0.05) (Table 4).

**Table 4.** Characteristics of *B. pertussis* isolates by age group, post-2020.

|  | ≤36 months | 37 months-18 years | >18years |
| --- | --- | --- | --- |
| Vaccine-Antigen Genotypes |  |  |  |
| *ptxC1/prn1/ptxP1/fhaB2400_5550-3/tcfA2* | 2 (8.70%) | 2 (2.78%) | 0 |
| *ptxC2/prn150/ptxP3/fhaB2400_5550-1/tcfA2* | 18 (78.26%) | 69 (95.83%) | 33 (100%) |
| *ptxC2/prn150/ptxP3/fhaB-2400_5550-1/tcfA9* | 1 (4.35%) | 0 | 0 |
| *ptxC2/prn2/ptxP3/fhaB-2400_5550-1/tcfA2* | 2 (8.70%) | 1 (1.39%) | 0 |
| MLVA types |  |  |  |
| MT28 | 14 (60.87%) | 38 (52.78%) | 27 (81.82%) |
| Others | 9 (39.13%) | 34 (47.22%) | 6 (18.18%) |
| Macrolide Resistance |  |  |  |
| S | 2 (8.70%) | 1 (1.39%) | 0 (100%) |
| R | 21 (91.30%) | 71 (98.61%) | 33 (0%) |

MT28 emerged as the dominant lineage across all age strata post-2020; with all isolates exhibiting macrolide-resistant and carrying the *ptxP3* allele (hereafter referred to as MT28-*ptxP3*-MRBP). Overall, this lineage accounted for 61.72% (79/128) of post-2020 isolates, with a significantly higher prevalence in adults (≥19 years) at 81.82% (27/33) than in children and adolescents (37 months–18 years) at 52.78% (38/72) (χ^2^ = 8.092, p < 0.01) (Table 4). No significant difference was observed between the ≥19 years group and infants aged ≤36 months (60.87%; p = 0.082). When infants and school‐age children/adolescents were combined into a single <19-year group and compared with the ≥19-year group, the age‐related difference in MT28-*ptxP3*-MRBP prevalence remained significant (p < 0.01).

### Phylogenetic analysis

Phylogenetic analysis of 178 *B. pertussis* isolates revealed two dinstinct clades defined by *ptxP1* and *ptxP3* (Fig. 2). Within the *ptxP3* clade, green-highlighted branches denote isolates carrying the A2047G resistance mutation, which closely coincides with the *prn150* genotype. No significant age‐related clustering was observed. The phylogeny of 615 Chinese *B. pertussis* isolates showed no provincial-level clustering, with Shanghai isolates genetically similar to the national population (Fig. 3). The MT28-*ptxP3*-MRBP lineage was first detected in Beijing in 2019 (SRR27796581, SRR27796588) and 2020 (SRR27796580), with the earliest Shanghai isolate of this lineage also in 2020 (SRR27796580). Global phylogenetic tree (Fig. 4) reveals strong geographic clustering of Chinese isolates. Outside China, only four MT28-*ptxP3*-MRBP isolates have been identified—ERR13476619 (2024, France), DRR631445 (2024, Japan), SRR32181461 (2024, USA), and SRR32181462 (2024, USA). Among all 1774 *B. pertussis* genomes analyzed, MT60 represents 1.41% (25/1,774), all post-2020; 22 of these (88%, 22/25) derive from this study, and the remaining three from Zhejiang Province, China.

**Fig. 2.**
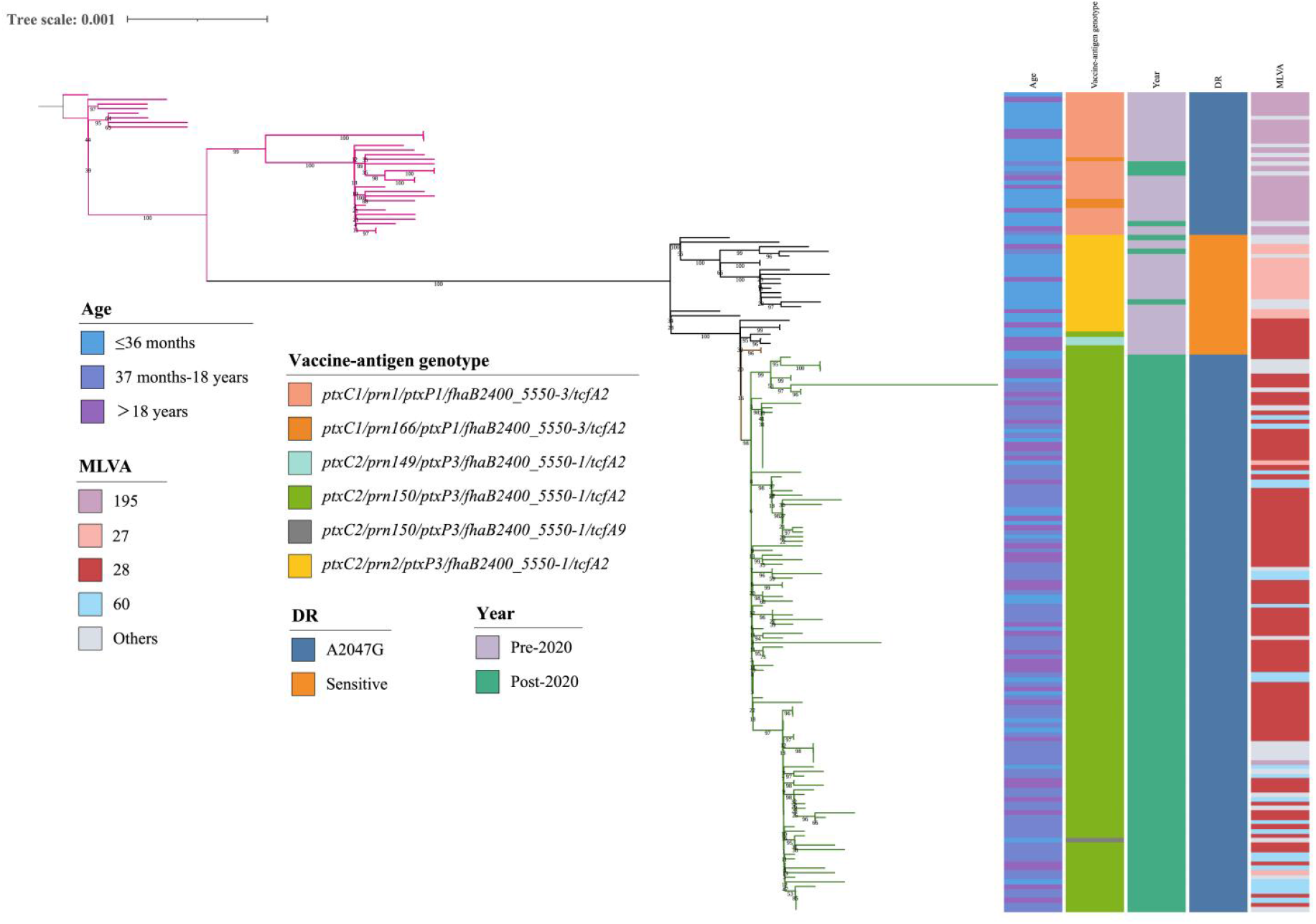
Maximum-likelihood phylogeny of 178 *B. pertussis* isolates from Shanghai (2018-2024). Branches are colored by *ptxP* allele clade: *ptxP1*(magenta) and macrolide-resistant *ptxP3* (green).

**Fig. 3.**
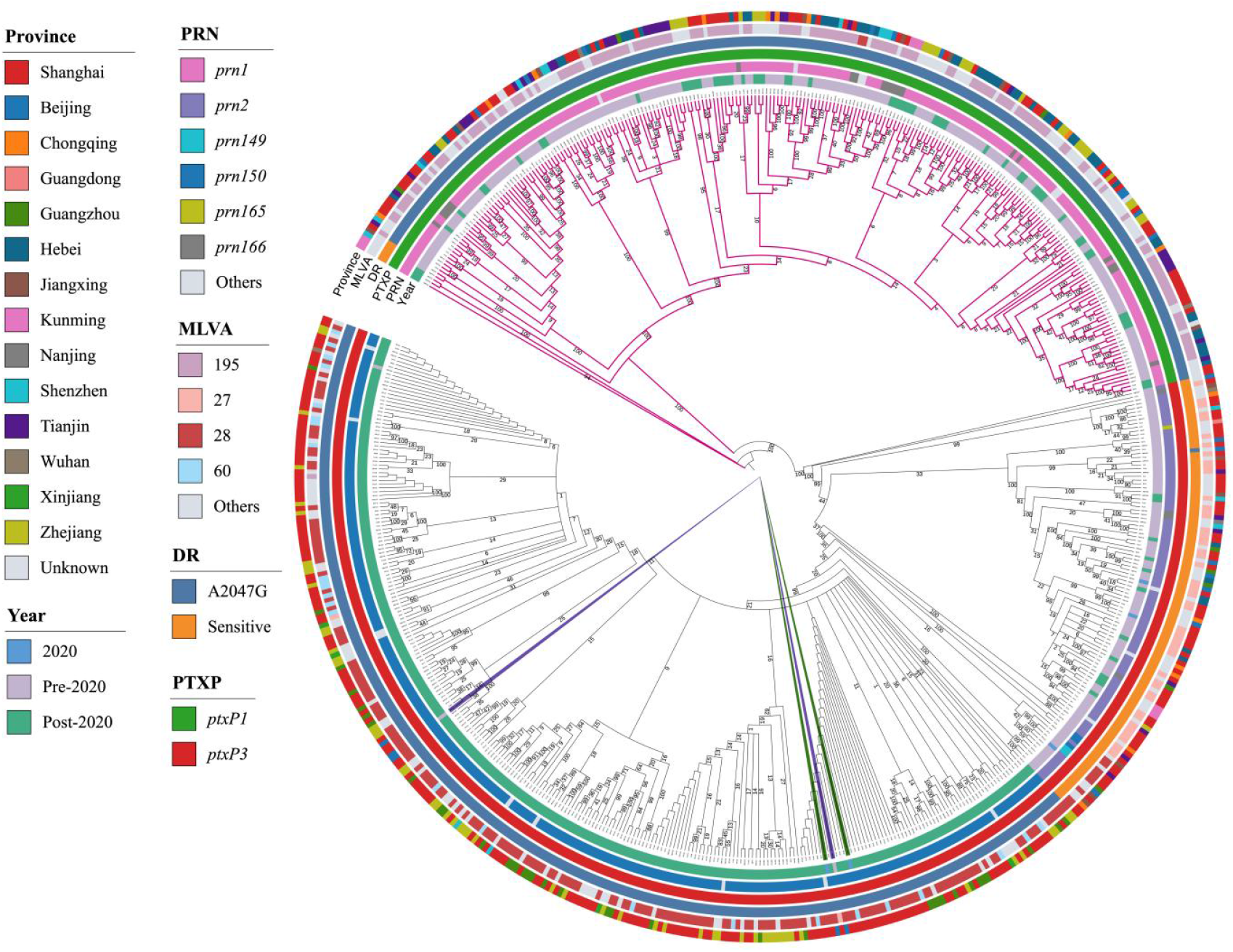
Phylogeny of 615 Chinese *B. pertussis* genomes. Branches belonging to the *ptxP1* clade are shown in magenta. MT28-*ptxP3*-MRBP isolates are highlighted in blue for those collected pre-2020 and in green for those collected in 2020. No clear clustering by province was observed among Chinese isolates.

**Fig. 4.**
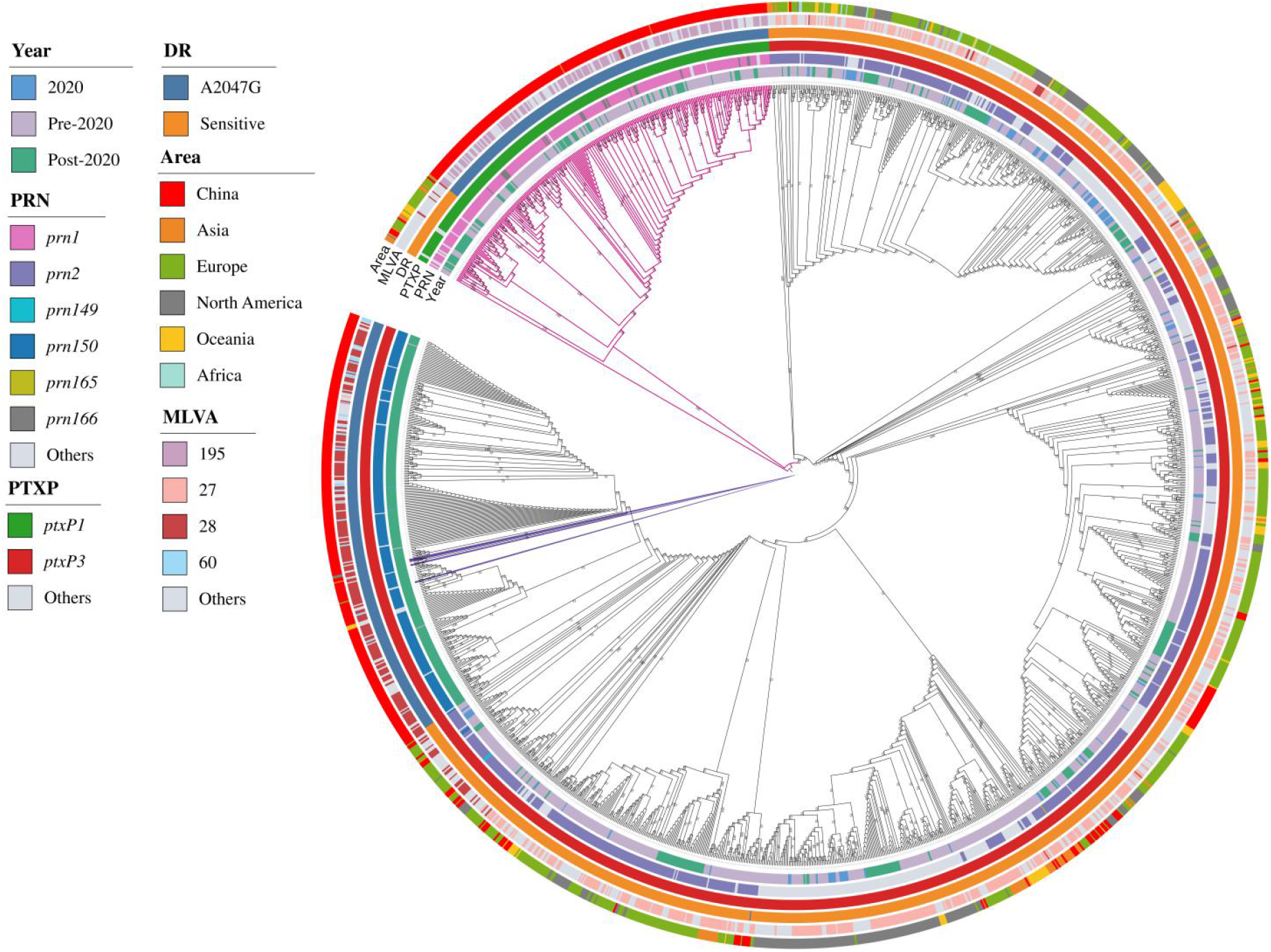
Phylogeny of 1774 global *B. pertussis* isolates. Branches colored in red represent the non-*ptxP3* clade. MT28-*ptxP3*-MRBP isolates from outside China are marked in purple.

## Discussion

In this study, we systematically analyzed the epidemiological characteristics, antimicrobial susceptibility, and genomic profiles of 178 *B. pertussis* isolates collected in Shanghai from patients of all age groups between 2018 and 2024. Our findings indicate that, in the pre-2020 period, pertussis cases occurred predominantly in infants aged ≤ 36 months, whereas in the post-2020 period they were mainly observed in school-age children and adolescents aged 37 months-18 years. The MRBP positivity rate rose from 58.33% in 2018 to 94.29% in 2021, and the MT28 and *ptxA1/ptxC2/prn150/ptxP3/fhaB2400_5550-1/fim2-1/tcfA2/fim3-1* genotypes rapidly became dominant post-2020. Importantly, this study is the first to demonstrate age-specific differences in the prevalence of the MT28-*ptxP3*-MRBP lineage in Shanghai after 2020.

In 2011, the first case of MRBP in China was reported in Shandong Province^[18]^. Li et al. conducted a multicenter study across northern and southern China from 2014 to 2016 and found that the MRBP positivity rate reached 91.1% (194/213) in northern isolates versus 64.3% (36/56) in southern isolates^[19]^. Fu et al. reported a 57.5% (81/141) MRBP rate in Shanghai during 2016-2017^[20]^. However, in the post-2020 period, MRBP positivity exceeded 97% in both northern and southern regions^[11, 12, 21]^. In our study, MRBP positivity remained at 46% pre-2020 and rose to 97.66% post-2020, consistent with the overall southern data and the findings of Fu et al.^[11]^ at Fudan Pediatric Hospital, Shanghai. Despite the high MRBP rates in China, much lower proportions have been reported elsewhere. For example, studies in France (June 2023-May 2024) and Finland (April-October 2024) found MRBP rates of only 1.5% (1/67) and 0.22% (1/462), respectively^[22, 23]^. Among the 1159 non-Chinese *B. pertussis* genomes included in our analysis, only nine (0.78%, 9/1159) harbored the 23S rRNA A2047G mutation. Nevertheless, this does not imply that MRBP is confined to China. For instance, although India accounts for 26.5% of global pertussis cases^[24]^, relatively few isolates have been characterized and data on antimicrobial susceptibility or A2047G mutation prevalence remain scarce^[25]^, leaving the true global burden of MRBP uncertain.

Currently approved aP vaccines contain up to five bacterial antigens: Pertussis Toxin (PTX) and four adhesion proteins, including Filamentous Hemagglutinin (FHA), Pertactin (PRN), and Fimbriae Types 2 and 3 (FIM2/3)^[26]^. The genotype of the Chinese *B. pertussis* vaccine strain is *ptxA2/ptxC1/prn1/ptxP1/fhaB2400_5550-1/fim2-1/tcfA2/fim3-1*^[27]^. The approved aP vaccines in China are mainly divided into two types: one containing PTX and FHA (two-component vaccine), and the other containing PTX, FHA, and PRN (three-component vaccine)^[28]^. This also partially explains why, in this study, all strains, except for one identified as a new *tcfA9* variant, were consistent with the vaccine strain, featuring *fim2-1/tcfA2/fim3-1*.

Two studies conducted in Beijing, China, reported that PRN in the region is predominantly *prn2*, with no detection of *prn150* (0/288 and 0/60)^[12, 29]^. However, in a study by Zhou et al.^[27]^ also conducted in Beijing, 100% (44/44) of *B. pertussis* isolates exhibited *prn150*, consistent with the findings in this study and the overall trend observed in Shanghai^[11]^. Both *prn150* and *prn2* are genetically distinct from the vaccine strain, and prn-deficient strains are widely present globally^[21, 27, 30, 31]^. FIM and FHA play crucial roles in allowing *B. pertussis* to evade immune surveillance during infection and to establish colonization in the respiratory tract^[32]^. Moreover, a mouse model demonstrated that mutants lacking FHA and FIM showed significantly reduced infectivity in the nasal cavity^[33]^. In this study, the proportion of *fhaB2400_5550-1* (vaccine strain genotype) increased from 46% (23/50) pre-2020 to 96.88% post-2020. Although this shift was unexpected, similar reports have emerged in several other studies in China^[27, 28]^. Among the 615 Chinese *B. pertussis* strains included in the analysis, the proportions of *fhaB2400_5550-1* were 34.42% (95/276) pre-2020 and 83.23% (278/334) post-2020.

Pre-2020, *B. pertussis* strains carrying the *ptxP3* allele had been reported in China, but *ptxP3*-MRBP was very rare, with *ptxP1*-MRBP being predominant^[13, 19, 34, 35]^. In this study, all 23 *ptxP3*-positive *B. pertussis* isolates from the pre-2020 period were macrolide-susceptible, while all 27 *ptxP1*-positive isolates were resistant. Post-2020, pertussis cases in China surged, accompanied by rapid expansion of *ptxP3*-MRBP, which became the dominant lineage^[12, 29, 36]^, consistent with the overall trends observed in this study. Additionally, this study found that MT28-*ptxP3*-MRBP was not first detected in Shanghai; it was identified in Beijing as early as 2019. Phylogenetic analysis of 1159 international *B. pertussis* isolates included in this study revealed that *ptxP3* was predominant globally even before 2020. Notably, MT28-*ptxP3*-MRBP was first detected in 2024 in Japan, France, and the USA, spanning three continents, indicating a potential for cross-border spread. Miettinen et al.^[22]^ also detected one *ptxP3*-MRBP isolate in a study of 462 *B. pertussis* isolates collected from different regions of Finland between April and October 2024, though MLVA typing data was not provided. The global phylogenetic tree reveals that Chinese *B. pertussis* strains exhibit strong geographic clustering, with MT28-*ptxP3*-MRBP emerging as a dominant lineage in China from 2019 to 2021 in a remarkably short period, which may indicate a high risk of international spread following its emergence.

In this study, we first observed significant differences in the prevalence of MT28-*ptxP3*-MRBP across different age groups after 2020. The proportion of MT28-*ptxP3*-MRBP was significantly higher in the ≥19 years group compared to the 37 months–18 years group. No significant difference was observed between the ≥19 years group and the ≤36 months group, possibly due to the smaller sample size in the latter group. However, a significant difference remained between the ≥19 years group (60.87%, 27/33) and the <19 years group (54.74%, 52/95), suggesting that MT28-ptxP3-MRBP may have a higher transmission advantage in older age groups.

In this study, MT60, a new genotype emerging post-2020, accounted for 17.19% (22/128) and was the second most common genotype after MT28. However, aside from the samples in this study, only three (0.19%, 3/1596) *B. pertussis* isolates from Zhejiang Province, China, displayed the MT60 genotype. All MT60 isolates were *ptxP3*-MRBP, and they showed high homology with MT28-*ptxP3*-MRBP in the phylogenetic tree, indicating that this lineage should be closely monitored.

There are some limitations to this study: first, the epidemiological information in the downloaded data was relatively limited, and the sample size was small, all collected from Shanghai, which may not be fully representative of the broader geographic distribution. Second, the isolation and culture of B. pertussis are challenging, and there may be some bias in the selection of isolates.

## Conclusion

In summary, this study is the first to identify significant differences in the prevalence of MT28-*ptxP3*-MRBP across age groups, suggesting that this *B. pertussis* genotype may have a transmission advantage in older populations. Moreover, MT28-*ptxP3*-MRBP has begun to appear in countries outside of China, indicating a risk of international spread. Therefore, we recommend further strengthening active surveillance across all age groups and closely monitoring the global spread of MT28-*ptxP3*-MRBP to inform and optimize pertussis control strategies.

## CRediT authorship contribution statement

**Zhen Xu**:Conceptualization, Data curation, Formal analysis, Writing – original draft, Investigation. **Zhuoying Huang**: Investigation, Methodology, Resources, Writing – original draft. **Lingyue Yuang**: Investigation, Methodology, Validation. **Huanyu Wu**: Software, Resources. **Xin Chen**: Visualization,Validation. **Min Chen**: Funding acquisition, Resources.**Yuan Zhuang**: Writing – review & editing, Supervision, Project administration, Funding acquisition, Resources. **Jun Feng**: Writing – review & editing, Supervision, Project administration, Funding acquisition, Resources..

## Data Availability

Data will be made available on request.

